# Relish/NF-κB acts in intestinal stem cells to promote epithelial repair in *Drosophila*

**DOI:** 10.1101/2022.09.29.510182

**Authors:** Meghan Ferguson, Minjeong Shin, Edan Foley

## Abstract

Immune signals coordinate the repair of damaged epithelia by intestinal stem cells. However, it is unclear if immune pathways act autonomously within the stem cell to direct the damage response pathway. We consider this an important question, as stem cell dynamics are essential for formation and maintenance of the entire epithelium. We used *Drosophila* to determine the impact of stem cell-specific loss of NF-κB on tissue regeneration upon chemical injury. We found that loss of NF-κB enhanced cell death, impaired enterocyte renewal and increased mortality. Mechanistically, we showed that the Ras/ERK pathway is essential for NF-κB-dependent maintenance of cell viability and tissue repair. Combined, our data demonstrate that stem cell-intrinsic NF-κB activity is essential for an orderly repair of damaged intestinal epithelia.

## INTRODUCTION

Intestinal Stem Cells (ISCs) generate the entire gut epithelium, a community of specialist cells that act in concert to extract nutrients, present a physical barrier to noxious agents, and protect the host from gut-resident microbes. Immune pathways are particularly important modifiers of epithelial responses to extrinsic stressors, and defective immune signals are linked to inflammatory illnesses alongside elevated risk of colorectal cancer (1, 2). For example, aberrant Tumor Necrosis Factor (TNF) signaling is a hallmark of inflammatory bowel disease, and TNF inhibitors are central to management of disease symptoms (3, 4). Despite the importance of immune activity for gut health, we know little about ISC-intrinsic requirements for immune signals to protect from acute epithelial insults. As ISCs are critical for formation and maintenance of the epithelium, we consider it important to resolve the extent to which immune activity directly impacts ISC function.

Numerous studies documented effects of immune signals from gut-resident leukocytes or stromal tissue on ISC proliferation and differentiation (5, 6), and aberrant immune activity is associated with intestinal inflammation and cancer(1, 7, 8). However, the cellular complexity of mammalian intestines, combined with a paucity of genetic reagents, has hampered our ability to identify ISC-intrinsic functions of immune signaling in vertebrates. In this context, *Drosophila melanogaster* is an excellent model to characterize genetic regulation of intestinal epithelial function(9). Like vertebrates, the fly gut is lined by a continuous epithelial layer that is maintained by basal, multipotent ISCs (10, 11). In flies and vertebrates, ISCs typically divide asymmetrically to generate a niche-resident daughter stem cell, and a transient, undifferentiated cell type that exits the niche and migrates apically, where it matures as an absorptive enterocyte, or a secretory enteroendocrine cell (12–15). In flies, the transient enterocyte precursor is classified as an enteroblast, and the stem cell-enteroblast pair constitutes the midgut progenitor compartment. Aside from similarities in cell identity and fate choices, ISC function is governed by similar signaling cassettes in flies and vertebrates. In both cases, cues from EGF, JAK-STAT and BMP pathways control ISC proliferation, while signals from the Notch pathway promote enterocyte fate choices among progenitors (12, 16–24). Given the extensive similarities between fly and vertebrate intestinal epithelial organization, *Drosophila* has considerable potential to uncover foundational aspects of immune regulation of epithelial stress responses.

The *Drosophila* Immune Deficiency (IMD) pathway, an evolutionary relative of TNF-receptor signaling, is a prominent regulator of fly responses to gut microbes (25, 26). Bacterial diaminopimelic-acid-containing peptidoglycan engages host pattern recognition proteins that signal through Imd to activate fly IKK and NF-κB orthologs (27). Active NF-κB modifies intestinal expression of antimicrobial peptides, stress response pathway genes, and genes that control metabolic activity (25, 26, 28–30). Recent work uncovered cell-type specific roles for the IMD-NF-κB axis in epithelial homeostasis, including regulation of ISC proliferation and differentiation (31, 32). In contrast, we know little about ISC-intrinsic roles for IMD in the control of stem cell responses to acute stresses.

We used the fly to characterize NF-κB-dependent control of ISC responses to acute epithelial stress. We discovered that damage response pathways activate NF-κB in stem cells, leading to Ras-dependent promotion of ISC survival and enhanced generation of transient enteroblasts. Targeted inactivation of NF-κB in ISCs resulted in widespread death of damaged stem cells, a failure to produce enterocyte precursors, and substantially impaired survival of challenged animals. Our work expands our appreciation of IMD-mediated control of gut homeostasis and uncovers a stem cell-intrinsic requirement for an NF-κB-Ras damage response in maintenance of the gut progenitor compartment.

## RESULTS

### NF-κB Regulates Intestinal Stem Cell Proliferation

Inflammatory signals disrupt intestinal epithelial growth, elevating the risk of chronic disease and colorectal cancer as the host ages. However, the intestine contains an intricate community of specialist cells, impairing our ability to systematically define lineage-specific contributions of immune activity to gut homeostasis. We depleted the NF-κB transcription factor family member *relish* (*rel*) exclusively from the progenitor compartment of adult fly intestines (*esg*^*ts*^*/rel*^*RNAi*^) and compared gut physiology to wild-type intestines (*esg*^*ts*^*/+*) raised under identical conditions. At early stages of adulthood, we noticed minimal differences between *esg*^*ts*^*/+* and *esg*^*ts*^*/rel*^*RNAi*^ flies. In both genotypes, the gut consisted of a pseudo-stratified epithelium, with evenly spaced GFP-marked progenitors interspersed among a regular lattice of mature enterocytes and enteroendocrine cells (Fig. 1A), as well as approximately equal incidences of ISC proliferation (Fig. 1C, day 10). Older wild-type intestines displayed classical hallmarks of age-linked dysplasia, including irregular epithelial structure (Figure 1B), and significantly elevated numbers of proliferative stem cells (Fig. 1C, day 30). In contrast, progenitor-restricted loss of *rel* prevented age-dependent decline of epithelial architecture (Fig. 1B) and resulted in significantly fewer stem cell mitoses (Fig. 1C, d30). Importantly, progenitor-specific knockdown of the IKKγ ortholog *kenny* (*key*) also prevented age-dependent increases in mitotic stem cells (Fig. 1D), confirming a central role for the IKK/NF-κB axis in age-associated deterioration of ISC function.

**Figure 1.**
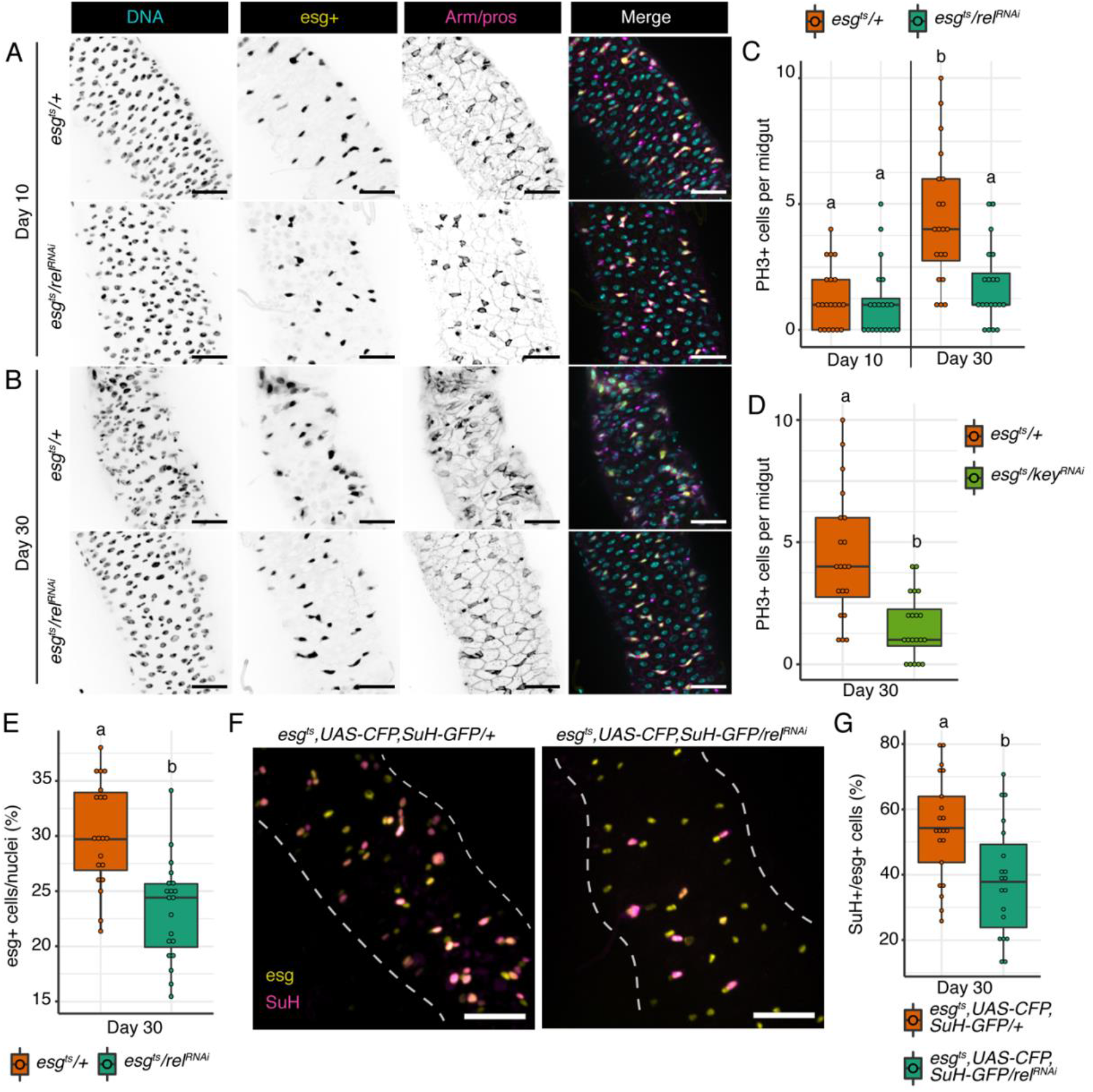
NF-κB regulates intestinal stem cell proliferation. **(A)** Posterior midgut of flies after 10 days of progenitor-specific NF-κB depletion (*esg*^*ts*^*/rel*^*RNAi*^) compared to wild-type (*esg*^*ts*^*/+)*. DNA labelled with Hoechst (cyan), progenitors labelled with esg (yellow), enteroendocrine cells labeled by pros (magenta) and cell borders labelled by Armadillo (magenta). **(B)** Posterior midgut of flies after 30 days of progenitor-specific *rel* depletion compared to wild-type. **(C)** PH3+ mitotic cells per midgut of wild-type and progenitor-specific *rel* depleted flies after 10 and 30 days. Significance found using ANOVA followed by pairwise Tukey tests. **(D)** PH3+ cells per midgut after progenitor-specific knockdown of IKKγ homolog *kenny* (*key*) **(E)** Proportion of esg+ progenitors per nuclei in 30 day old flies upon progenitor-specific *rel* knockdown. **(F)** Images from *esg*^*ts*^, *UAS-CFP, Su(H)-GFP* flies after 30 days of progenitor-specific *rel* depletion. Progenitors labelled with esg (yellow) and enteroblasts labelled with Su(H) (magenta). **(G)** Proportion of Su(H)+ enteroblasts within the progenitor pool upon progenitor-specific NF-κB knockdown. For D,E and G significance found using Student’s t test. Different letters denote significance at p < 0.05. Scale bars for A, B and F are 25μm.

As stem cell divisions maintain the progenitor population that generates a mature epithelium, we quantified the impacts of *rel* depletion on the number and identity of midgut progenitors in older flies. Consistent with a link between progenitor immune activity and ISC proliferation (Figure 1C-D), we observed significantly fewer progenitors in posterior midguts of *esg*^*ts*^*/rel*^*RNAi*^ flies relative to *esg*^*ts*^*/+* counterparts (Fig. 1E). To test if NF-κB activity also influences cell identity within the progenitor compartment, we determined the stem cell to enteroblast ratio in midguts of *esg*^*ts*^*/rel*^*RNAi*^ flies and *esg*^*ts*^*/+* controls. To do so, we employed a line that allows progenitor-specific depletion of *rel* while marking enteroblasts with CFP and GFP reporters, and ISCs with CFP alone (*esgGAL4 UAS-CFP, Su(H):GFP/UAS-rel*^*RNAi*^; *GAL80*^*ts*^, Fig. 1F). In wild-type intestines (*esgGAL4 UAS-CFP, Su(H):GFP/+; GAL80*^*ts*^), roughly 50% of all progenitors expressed enteroblast markers, suggesting approximately equal numbers of stem cells and enteroblasts (Figure 1G). In contrast, only 40% of *rel-*deficient progenitors expressed enteroblast markers, indicating that loss of NF-κB impaired enteroblast generation. Combined, our data implicate intestinal NF-κB activity in the proliferation and identity of midgut progenitors.

### ISC-specific NF-κB restricts proliferation upon damage

As our results imply that progenitor-specific NF-κB activity alters ISC proliferation, we determined the consequences of compromised progenitor cell immunity for epithelial responses to acute damage. Specifically, we characterized progenitor dynamics in *esg*^*ts*^*/rel*^*RNAi*^ and control *esg*^*ts*^*/+* flies that we fed dextran sodium sulfate (DSS), a toxic polysaccharide that disrupts the epithelium, promoting a compensatory burst of ISC divisions (19, 33–36). Consistent with Figure 1, progenitor-restricted inactivation of *rel* had no visible effect on intestinal physiology in unchallenged, ten-day-old flies (Fig. 2A). DSS caused extensive damage to *esg*^*ts*^*/+* midguts (Fig. 2B), leading to extra divisions (Fig. 2C) that increased ISC and progenitor numbers throughout the posterior midgut (Fig. S1A-B). We observed similar amounts of epithelial damage (Fig. 2B), and expansion of ISC and progenitor populations (Fig. S1A-B) in DSS-treated *esg*^*ts*^*/rel*^*RNAi*^ guts. However, we also uncovered a distinct effect of *rel* inactivation on damage-dependent proliferation. Specifically, we found that depletion of *rel* from progenitors nearly doubled the mitotic activity of ISCs in flies challenged with DSS compared to wild-type controls (Fig. 2C). Notably, progenitor-specific loss of *rel* also significantly increased the frequency of ISC proliferation in flies orally challenged with the enteric pathogen *Ecc15* (Fig. 2D), indicating a general requirement for NF-κB to inhibit ISC proliferation after ingestion of harmful agents.

**Figure 2.**
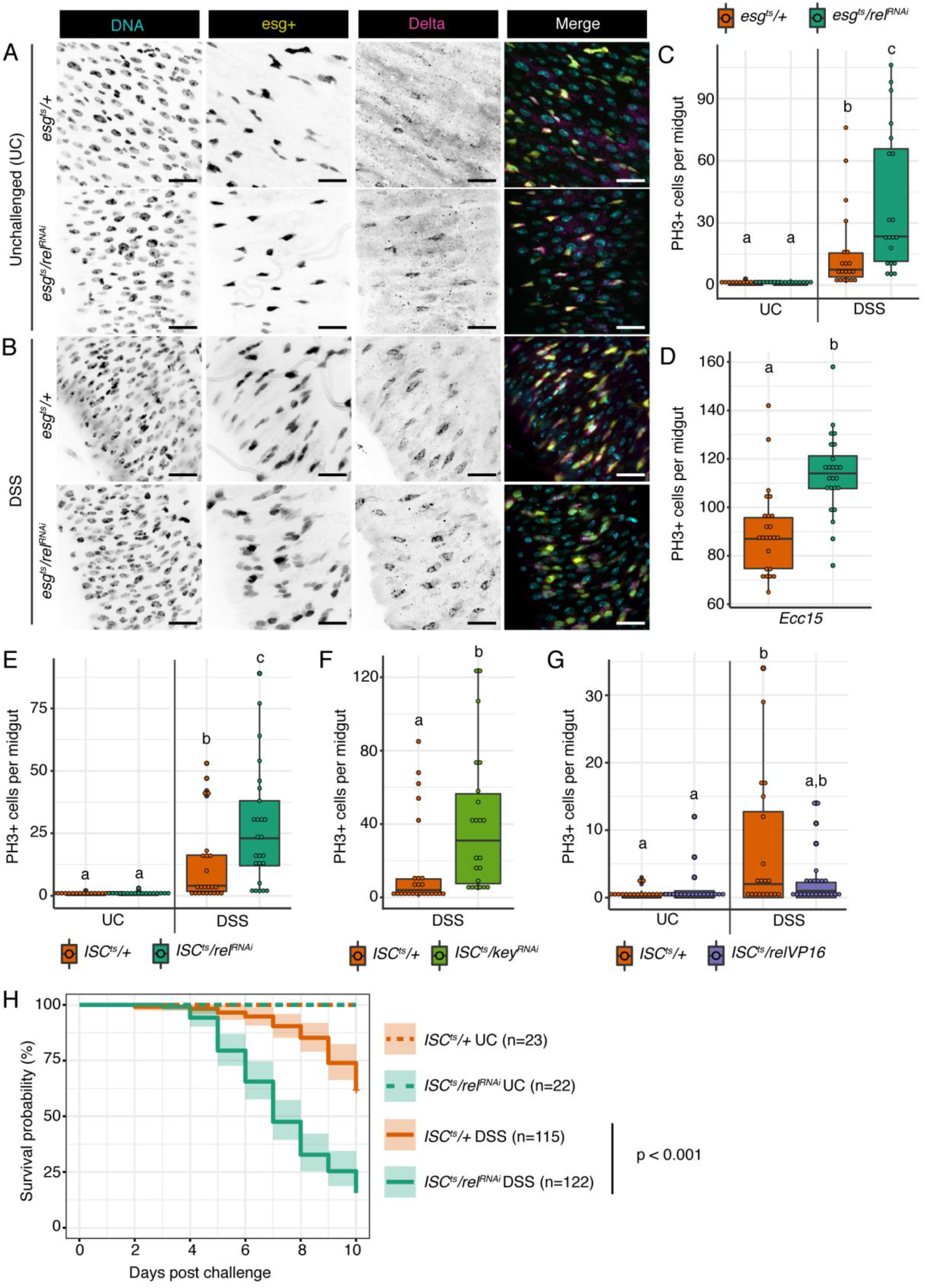
ISC-specific NF-κB restricts proliferation upon damage. **(A)** Posterior midgut of 10 day old unchallenged (UC) *esg*^*ts*^*/+* and *esg*^*ts*^*/rel*^*RNAi*^ flies fed PBS/5% sucrose solution for 48hrs. **(B)** Posterior midgut of 10 day old *esg*^*ts*^*/+* and *esg*^*ts*^*/rel*^*RNAi*^ flies fed 3% DSS/PBS/sucrose solution for 48hrs. Scale bars = 25μm. DNA labelled by Hoechst (cyan), esg+ progenitors (yellow), ISCs labelled with Delta (magenta). **(C)** PH3+ cells per midgut of 48hr UC and DSS treated *esg*^*ts*^*/+* and *esg*^*ts*^*/rel*^*RNAi*^ flies. **(D)** PH3+ cells per midgut of *esg*^*ts*^*/+* and *esg*^*ts*^*/rel*^*RNAi*^ flies fed *Ecc15* for 48hrs. **(E)** PH3+ cells per midgut of UC and DSS treated *ISC*^*ts*^*/+* and *ISC*^*ts*^*/rel*^*RNAi*^ flies. **(F)** PH3+ cells per midgut of UC and DSS treated *ISC*^*ts*^*/+* and *ISC*^*ts*^*/key*^*RNAi*^ flies. **(G)** PH3+ cells per midgut of UC and DSS treated *ISC*^*ts*^*/+* and *ISC*^*ts*^*/relVP16* flies. **(H)** Survival of *ISC*^*ts*^*/+* and *ISC*^*ts*^*/Rel*^*RNAi*^ flies upon exposure to 10% DSS. Significance for H found using log rank test. Significance for C, E and G found using ANOVA followed by pairwise Tukey tests. Significance for D and F found using Students t test. Different letters denote significance at p < 0.05.

To identify the exact progenitor cell type where NF-κB acts to control damage-dependent growth, we knocked down *rel* exclusively in ISCs (*ISC*^*ts*^*/rel*^*RNAi*^) or enteroblasts (*Su(H)*^*ts*^*/rel*^*RNAi*^) and measured DSS-mediated ISC proliferation. Like progenitor-wide knockdown, ISC-specific loss of *rel* (Fig. 2E), or the IKKγ ortholog *key* (Fig. 2F) increased mitoses in response to DSS compared to wild-type controls, demonstrating an ISC-autonomous role for IKK/NF-κB activity in damage-responsive proliferation. Supporting a direct role for *rel* in the control of ISC proliferation, we also found that ISC-restricted expression of a constitutively active *rel* variant (*relVP16*) was sufficient to prevent DSS-responsive proliferation (Fig. 2G). In contrast, *rel* depletion from enteroblasts failed to increase ISC proliferation in DSS-treated flies (Fig. S1C), suggesting that *rel* acts primarily in ISCs to regulate damage-dependent proliferation.

Since rapid, orderly epithelial repair is essential to survive acute tissue damage, we measured the effect of ISC-specific loss of *rel* on survivability upon DSS exposure. Loss of *rel* did not affect the short (Fig. 2H), or long-term viability (Fig. S1D) of unchallenged flies. However, depletion of *rel* from ISCs significantly impaired the ability of flies to survive ingestion of DSS (Fig. 2H), confirming an essential role for Rel in ISC responses to damaging agents.

### ISC-specific NF-κB is required for enterocyte renewal upon damage

To understand how NF-κB regulates stem cell proliferation in times of acute tissue damage, we performed single-cell gene expression analysis on midguts dissected from unchallenged, or DSS-treated, *ISC*^*ts*^*/+* and *ISC*^*ts*^*/rel*^*RNAi*^ flies. We selected the *ISC*^*ts*^ driver line for this experiment, as it permits inactivation of *rel* exclusively in ISCs, while marking ISCs with YFP. As a result, we were able to resolve the impacts of ISC-restricted *rel* inactivation on all intestinal cell types, including the stem cell. In agreement with Figures 1 and 2, transcriptional states within unchallenged ten-day-old *ISC*^*ts*^*/+* and *ISC*^*ts*^*/rel*^*RNAi*^ midguts were broadly similar. In both instances, we identified approximately equal ratios of progenitor, enteroendocrine, and enterocyte lineages, as well as specialist copper and large flat cells (Fig. 3A, Fig. S2A). We also discovered a transcriptionally distinct cell population that expressed classical progenitor markers such as *esg* and *E*(*spl*)*m3-HLH*, alongside enterocyte markers such as the *trypsin* family of proteases (Fig. S2B). In each case, marker expression was at an intermediary level between that seen in progenitors and enterocytes (Fig. 3B-C), suggesting that these cells represent a transition state between undifferentiated progenitors and mature enterocytes. Therefore, we have labeled these cells premature enterocytes (preECs).

**Figure 3.**
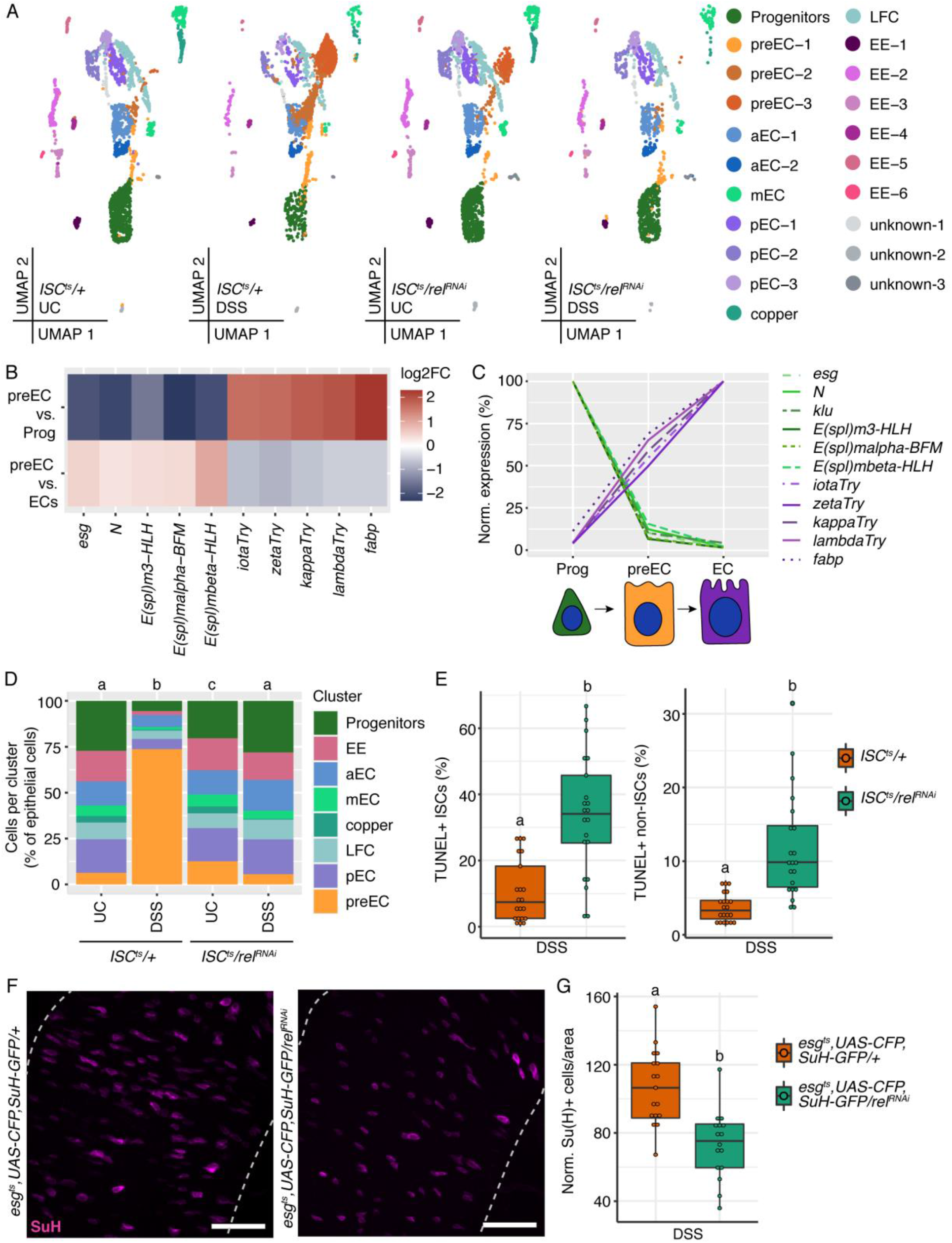
ISC-specific NF-κB is required for enterocyte renewal upon damage. **(A)** UMAP plots of UC and DSS treated *ISC*^*ts*^*/+* and *ISC*^*ts*^*/rel*^*RNAi*^ intestines from single-cell sequencing data. **(B)** Differential expression analysis comparing preECs to progenitors or enterocytes. All genes listed are p<0.05. **(C)** Normalized average expression values for select genes in progenitors, preECs and enterocytes. **(D)** Cells per cluster as a percent of total epithelial cells from UMAPs seen in A. Significance found using chi squared test followed by proportions test. **(E)** Proportion of ISCs and non-ISCs that are TUNEL positive in DSS treated *ISC*^*ts*^*/+* and *ISC*^*ts*^*/rel*^*RNAi*^ intestines. **(F)** Representative images of *esg*^*ts*^, *UAS-CFP, Su(H)-GFP* intestines upon DSS exposure and *rel* knockdown. SuH labels enteroblasts (magenta). Scale bars = 25μM. **(G)** Number of Su(H)+ enteroblasts per area upon DSS exposure and progenitor wide *rel* knockdown. Significance for E and G found using Students t test. Different letters denote significance at p <0.05.

In the absence of an extrinsic insult, preECs accounted for roughly 6% of all profiled cells in *ISC*^*ts*^*/+* midguts and 12% of all profiled cells in *ISC*^*ts*^*/rel*^*RNAi*^ midguts (Fig. 3D). Consistent with the extensive tissue renewal required to survive epithelial damage, ingestion of DSS caused a massive spike of preEC numbers in *ISC*^*ts*^*/+* flies. After 48h, 74% of all profiled cells in DSS-treated *ISC*^*ts*^*/+* guts expressed preEC markers, indicating accumulation of cells poised to replace dead and dying enterocytes (Fig. 3D). Notably, depletion of *rel* from stem cells resulted in an apparent failure to accumulate cells that expressed preECs markers. In marked contrast to midguts of DSS-treated *ISC*^*ts*^*/+* flies, only 6% of all profiled cells from DSS-treated *ISC*^*ts*^*/rel*^*RNAi*^ midguts expressed preEC markers (Fig. 3D), suggesting a possible failure of *rel*-deficient ISCs to generate preECs in response to damage.

Our initial results established that DSS caused a greater proliferative response in *rel*-deficient ISCs than wild-type counterparts (Fig. 2). However, our transcriptional data indicated an apparent absence of preECs, prompting us to ask if the hyperproliferation observed in DSS-challenged, *rel*-deficient ISCs productively contributes to epithelial renewal. To address this question, we measured cell death and enteroblast numbers in midguts of *ISC*^*ts*^*/+* and *ISC*^*ts*^*/rel*^*RNAi*^ flies that we challenged with DSS. To assess death, we stained DSS-treated *ISC*^*ts*^*/+* and *ISC*^*ts*^*/rel*^*RNAi*^ midguts with TUNEL and found that depletion of *rel* from ISCs significantly increased the amounts of TUNEL+ ISCs and epithelial cells (Fig. 3E), indicating that *rel* activity in ISCs maintains intestinal epithelial cell viability.

Under normal conditions, ingestion of DSS prompts the accumulation of enteroblasts as the gut initiates proliferative responses that replenish dying enterocytes (34). To quantify enteroblast numbers in response to damage we used the *esgGAL4 UAS-CFP, Su(H):GFP/UAS-relRNAi; GAL80*^*ts*^ line to mark enteroblasts with CFP and GFP while depleting *rel* from progenitors. We found that progenitor-specific loss of *rel* significantly reduced the number of enteroblasts upon DSS exposure when compared to wild-type controls (Fig. 3F, G). Taken together, our data show that NF-κB-deficient ISCs are more prone to cell death and are impaired in their ability to generate enteroblasts after exposure to DSS.

### ISC-specific NF-κB alters Hippo and EGF/Ras/MAPK pathway expression in response to damage

To resolve the mechanistic basis for NF-κB-dependent control of epithelial proliferative responses to damage, we compared the transcriptional responses of *ISC*^*ts*^*/+* and *ISC*^*ts*^*/rel*^*RNAi*^ midguts to DSS. First, we established cell-type specific transcriptional responses of a wild-type intestine to damage by identifying differentially expressed genes in an integrated data set generated from expression profiles of unchallenged and DSS-treated *ISC*^*ts*^*/+* midguts. As expected, DSS ingestion resulted in lineage-specific impacts on expression of numerous genes involved in growth, differentiation, and cell migration (Fig. S3A). In particular, progenitors responded to DSS with decreased expression of Notch targets and increased expression of EGF, Hippo, JNK, and JAK-STAT regulators, central elements of the proliferative epithelial repair pathway(37) (Fig. S3B). Notably, DSS treatment also elevated expression of the *rel* targets *pirk* and *PGRP-SC2* in progenitors (Fig. S3B), confirming that damage activates NF-κB in progenitors.

To determine how ISC-specific loss of NF-κB affects the gut response to damage, we then compared gene expression in DSS-treated *ISC*^*ts*^*/+* and *ISC*^*ts*^*/rel*^*RNAi*^ single-cell transcriptomes. Among progenitors, ISC-specific loss of *rel* significantly affected DSS-dependent expression of genes required for cell migration, differentiation, and stem cell proliferation (Fig. 4A). Conversely, loss of *rel* in ISCs primarily impacted DSS-dependent expression of genes linked with metabolism in enteroendocrine cells and enterocytes (Fig. 4A). This indicates that ISC-specific NF-κB knockdown primarily alters epithelial renewal in a cell-autonomous fashion, although we did find evidence that blocking *rel* in ISCs affects the Hippo pathway in copper cells (Fig. S4). A more detailed comparison of wild type and *rel*-deficient progenitor transcriptomes showed that NF-κB depletion decreased expression of *relish* and its target genes, confirming successful knockdown of *rel* (Fig. 4B). In addition to effects on immune response regulators, NF-κB knockdown altered expression of multiple Hippo and EGF/Ras regulators in progenitors, including the EGF inhibitor *sprouty (sty)*, the signaling regulator *Star* (*S*), and the EGF/Ras-responsive transcription factor *pointed* (*pnt*) (Fig. 4B), suggesting possible links between NF-κB and EGF/Ras signaling in progenitors.

**Figure 4.**
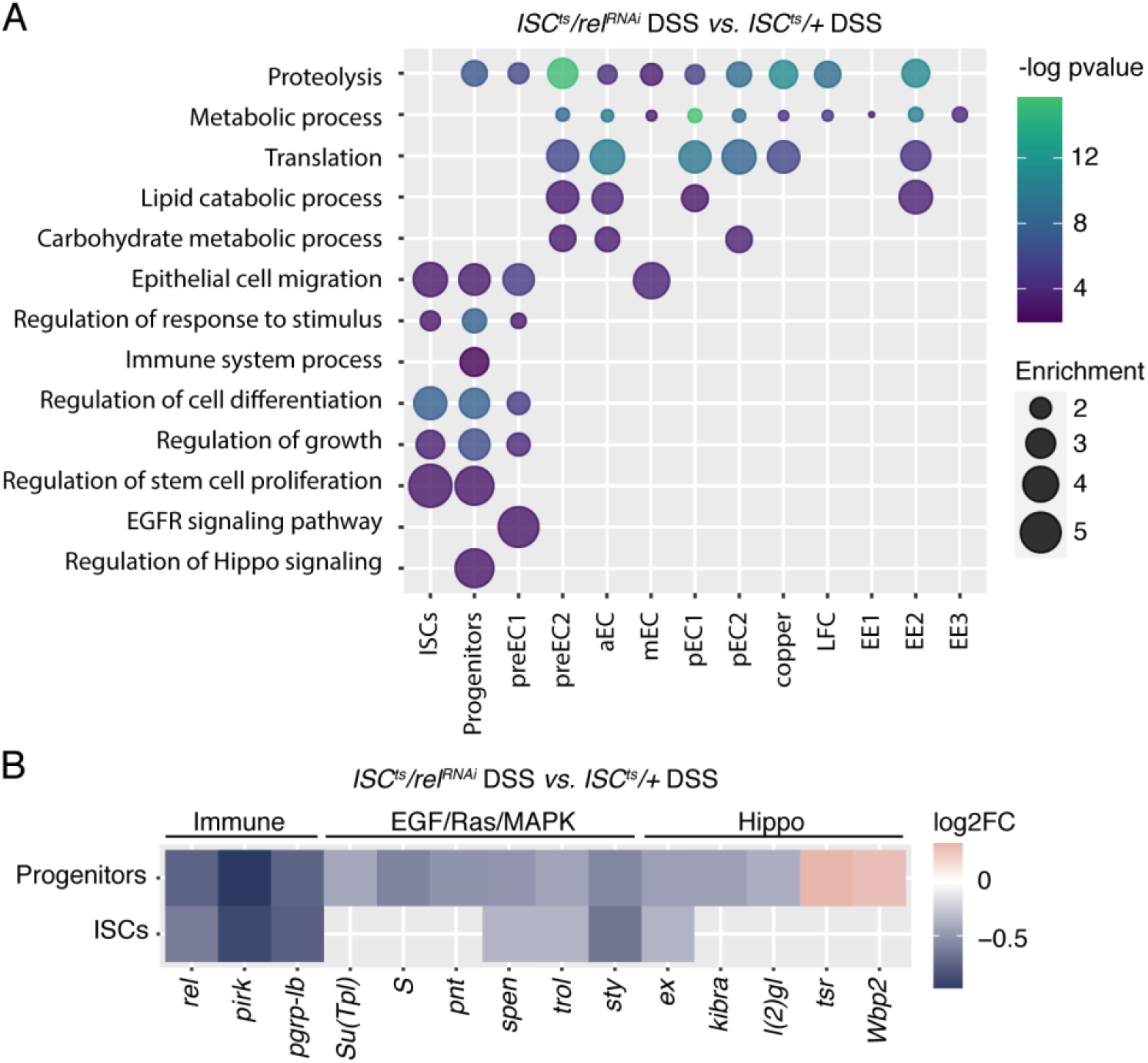
ISC-specific NF-κB alters Hippo and EGF/Ras pathway expression in response to damage. **(A)** GO term bubble plot of biological processes significantly enriched in DSS treated *ISC*^*ts*^*/rel*^*RNAi*^ intestines compared to DSS treated *ISC*^*ts*^*/+*. Size of the bubble shows GO term enrichment score and color shows −log pvalue. **(B)** Genes differentially expressed in Progenitors and EYFP+ Progenitors (ISCs) in DSS treated *ISC*^*ts*^*/rel*^*RNAi*^ intestines compared to DSS treated *ISC*^*ts*^*/+*. All genes shown are p < 0.05.

### Ras/ERK acts downstream of NF-κB in ISCs to regulate intestinal repair

EGF/Ras controls proliferation, differentiation, and survival in progenitors. Therefore, we asked if Ras acts downstream of NF-κB in the control of stem cell proliferation. To determine if NF-κB impacts Ras activity we first measured phosphorylated ERK (pERK) in DSS-treated *ISC*^*ts*^*/+* and *ISC*^*ts*^*/rel*^*RNAi*^ midguts (Fig. 5A,B). With DSS exposure ~40% of wild-type ISCs are pERK+, however, this decreases to ~5% upon ISC-specific *rel*-depletion (Fig. 5C), suggesting that NF-κB is necessary for Ras activation in ISCs upon damage. Next, we asked whether inhibition of Ras alone alters proliferation in response to DSS. Consistent with earlier reports (17, 38, 39), progenitor-wide inhibition of Ras (*RasN17/+;esg*^*ts*^*/+*) blocked mitosis upon DSS exposure (Fig. S5). However, inactivation of Ras in ISCs (*RasN17/+;ISC*^*ts*^*/+*, Fig. 5D) increased proliferation in response to DSS when compared to wild-type intestines (Fig. 5E), a phenotype similar to ISC-specific *rel* depletion. In addition, and similar to the consequences of *rel* depletion, ISC-specific inactivation of Ras increased the number of apoptotic cells in intestines of DSS-treated flies (Fig. 5F), suggesting that Ras may act downstream of Relish in ISCs to control epithelial viability and proliferation responses to DSS.

**Figure 5.**
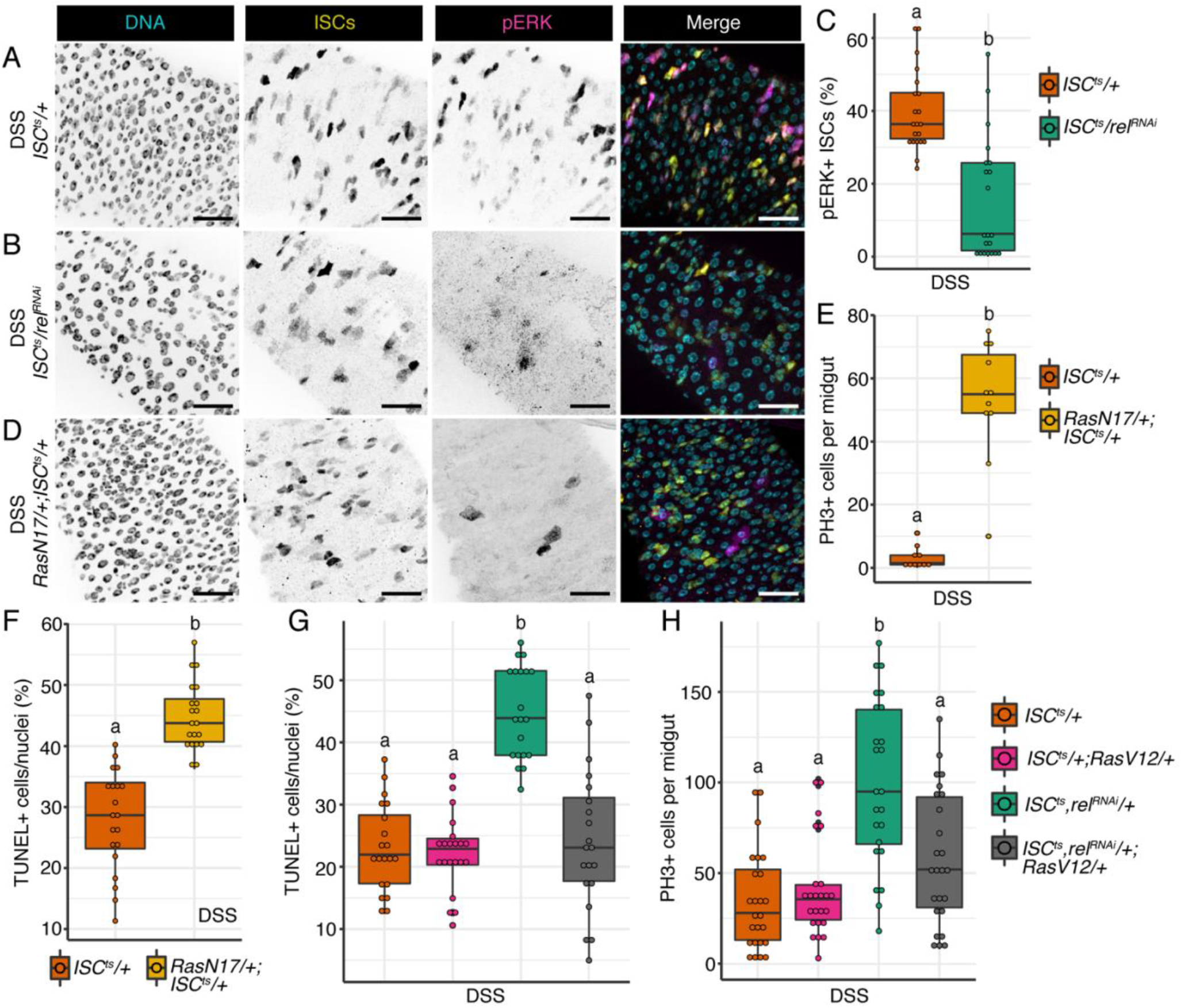
Ras/ERK acts downstream of NF-κB in ISCs to regulate intestinal repair. **(A)** Representative images of a wild-type *ISC*^*ts*^*/+* intestine exposed to DSS. DNA labelled with Hoechst (cyan), ISCs tagged with EYFP (yellow) and pERK labels Ras active cells (magenta). **(B)** Images of *ISC*^*ts*^*/rel*^*RNAi*^ intestine exposed to DSS. **(C)** Proportion of pERK+ ISCs upon DSS exposure and ISC-specific *rel* knockdown. **(D)** Images of *RasN17/+;ISC*^*ts*^*/+* where Ras is blocked specifically in ISCs. Scale bars for A, B and D are 25μm. **(E)** PH3+ cells per midgut upon DSS exposure and ISC-specific Ras inhibition. **(F)** Proportion of TUNEL+ nuclei in the posterior midgut upon ISC-specific Ras inhibition and DSS exposure. **(G)** TUNEL+ cells per nuclei in posterior midguts upon DSS challenge and ISC-specific *rel* knockdown (*ISC*^*ts*^,*rel*^*RNAi*^*/+)* and Ras activation (*ISC*^*ts*^*/+;RasV12/+)* alone or in conjunction (*ISC*^*ts*^,*rel*^*RNAi*^*/+;RasV12/+)*. **(H)** PH3+ cells per midgut upon DSS challenge and ISC-specific *rel* knockdown and Ras activation alone or in conjunction. Significance found for C E and F using Students t test. Significance for G-H found using ANOVA followed by multiple pairwise Tukey tests. Different letters denote significance at p <0.05.

Since *rel* depletion diminished Ras activation, and ISC-specific Ras inhibition phenocopies several aspects of Rel inactivation, we next asked if Ras acts downstream of NF-κB in the context of damage. To determine the relationship between NF-κB and Ras in ISCs we knocked down *rel* and activated Ras concurrently in ISCs of flies that we challenged with DSS (*ISC*^*ts*^,*rel*^*RNAi*^*/+;RasV12/+*). Ras activation in ISCs alone had no effect on the number of cells undergoing apoptosis or mitosis upon DSS exposure (Fig. 5G-H). Conversely, ISC-specific depletion of *rel* increased the numbers of apoptotic cells and promoted proliferation in response to DSS (Fig. 5G-H). Interestingly, when we activated Ras in *rel*-deficient ISCs, we found that the levels of apoptosis and mitosis decreased to wild-type levels (Fig. 5G-H). Together these results indicate that NF-κB acts in ISCs to promote cell survival by activating the Ras pathway. In the absence of Relish, Ras activity is diminished which induces apoptosis and provokes ISCs to divide excessively to compensate for the lack of effective repair.

## DISCUSSION

Intestinal stem cells adapt gut physiology in response to extrinsic immune signals. For instance, lymphoid and myeloid cells direct ISC proliferation to match the rate of epithelial damage caused by microbial challenges (5), while germline-encoded pattern recognition receptors, such as TLR4 and NOD2, act within the epithelium to modify ISC survival and proliferation during periods of acute, toxic stress (40, 41). Collectively, these observations emphasize the impact of immune activity on ISC survival and proliferation. However, the complexity of cell types within an animal gut makes it difficult to accurately resolve the impacts of ISC-intrinsic immunity on gut function. We consider this a particularly important question, as ISCs are essential for the formation of a functional, protective epithelium, and disruptions to the gut epithelium often lead to severe inflammatory disease.

To examine links between stem cell immunity and epithelial organization, we depleted the IKK/NF-κB orthologs *kenny* and *relish* from adult fly midgut ISCs and monitored epithelial maintenance during ageing, or during acute stress. In agreement with recent work (31), we found that ISC-specific inactivation of *rel* prevented the hyperplastic decay in epithelial organization associated with ageing, suggesting that ISC-intrinsic immune signals may contribute to loss of gut function in elderly flies. In addition to modifying homeostatic proliferation, previous studies suggested that the IMD pathway affects fate choices made by differentiating precursors. For instance, progenitor-specific inhibition of the Imd protein increased enteroblast numbers and diminished enteroendocrine cell numbers, suggesting that Imd promotes an enteroendocrine cell fate (31). In contrast, we discovered that depletion of *relish* from progenitors decreased enteroblast numbers. In combination, these data suggest that activation of Imd in the absence of a downstream NF-κB response promotes enterocyte fate choices, whereas engagement of an intact IMD pathway favors enteroendocrine cell development. In agreement, a recent study showed that knockdown of *relish* in progenitors increased enteroendocrine cell numbers in the midgut (32).

To understand how Relish controls epithelial repair, we combined imaging, genetics and single-cell transcriptomics to establish the ISC response to damage in the presence or absence of *rel*. We found that ISC-specific loss of *rel* resulted in extensive, stress-dependent ISC death that impaired damage-responsive generation of pre-enterocytes, and enhanced fly sensitivity to DSS. Our observations contrast with the role of Relish in enterocytes, where it contributes to cell death by orchestrating the lumenal expulsion of dying cells (42), indicating elaborate, cell-specific contributions of the IMD pathway to protection from damage.

Mechanistically, our data implicate Ras/ERK as a key mediator of Rel-dependent ISC viability. We discovered ISC-intrinsic impacts of Rel on expression of multiple EGF/Ras pathway elements, and we found that Relish is essential for DSS-dependent activation of ERK. Consistent with a link between Rel and Ras in ISC viability, ISC-specific inactivation of Ras mimics the apoptotic, hyperproliferative phenotype observed with ISC-specific inactivation of Rel. Additionally, we noticed that genetic activation of Ras exclusively in stem cells overrides the hyperproliferative phenotype associated with Rel-deficiency. From these results, we speculate that epithelial stress engages NF-κB in ISCs, leading to Ras/ERK activation, which promotes stem cell viability, attenuates stem cell proliferation and permits generation of adequate pre-enterocyte numbers to protect the interior from excess damage. Currently, it is unclear how Rel activates Ras/ERK, or whether links between cell viability and proliferation are correlative or causative. In the future, it will be interesting to identify the signals that connect NF-κB and Ras/ERK, and to explore the impacts of modified ISC viability on cell proliferation and pre-enterocyte generation during acute stress.

We also believe our results raise an interesting aspect of the role of the Ras/ERK pathway in fly midgut progenitors. Our data agree with earlier reports that progenitor-wide inhibition of the Ras-ERK pathway blocks the proliferative burst typically seen in midguts of flies exposed to noxious agents (17, 38, 39). However, we also found that inhibition of Ras exclusively in stem cells significantly increases the rate of damage-response proliferation. These observations raise the possibility that EGF/Ras alters proliferation via different mechanisms in ISCs and enteroblasts. We consider it possible that EGF/Ras acts in ISCs to support epithelial survival during damage, thereby reducing the proliferative requirement for ISCs. In support of this, inactivation of EGF in progenitors results in increased apoptosis in ISC progeny (43). Together these observations suggest that Relish acts through EGF/Ras in ISCs to control cell survival in the face of extrinsic insults and are key factors in effective epithelial repair. Given the constant environmental fluctuations the intestinal lumen is exposed to, we argue that immune signaling in ISCs is used as an adaptive mechanism to tune cell survival, differentiation and proliferation to the specific needs of the epithelium.

## ACKNOWLEDGEMENTS

We thank the Bloomington *Drosophila* Stock Center, Dr. Bruce Edgar and Dr. Lucy O’Brien for providing fly lines. We acknowledge microscopy support from Drs. Steven Ogg and Gregory Plummer at the Faculty of Medicine and Dentistry Imaging core. We acknowledge support from Drs. Joaquin Lopez-Orozco and Sudip Subedi the High Content Analysis core. This work was supported by a grant from the Canadian Institute of Health Research to EF (Grant # PJT 159604). M.F. has funding through Alberta Innovates Graduate Student Scholarships and NSERC PGS-D. M.S. has funding through Basic Science Research Program through the National Research Foundation of Korea (NRF) funded by the Ministry of Education (2020R1A6A3A0303955511).

## MATERIALS AND METHODS

### *Drosophila* husbandry

*Drosophila* crosses were setup and maintained at 18°C on standard corn meal food (Nutri-Fly Bloomington formulation; Genesse Scientific). All experimental flies were virgin female and kept at a 12h:12h light dark cycle throughout. Upon eclosion, flies were kept at 18°C then shifted to the appropriate temperature once 25-30 flies per vial was obtained. Fly lines used in this study were: *w;esg-GAL4,tubGAL80*^*ts*^,*UAS-GFP* (*esg*^*ts*^) (Bruce Edgar), *w;esg-GAL4,UAS-2xEYFP;Su(H)GBE-GAL80,tub-GAL80*^*ts*^ (*ISC*^*ts*^) (Bruce Edgar), *w;Su(H)GBE-GAL4,UAS-GFP;ubi-GAL80*^*ts*^ (*SuH*^*ts*^) (Bruce Edgar), *w*^*1118*^ (VDRC #60000), *relRNAi* (VDRC #49413), *keyRNAi* (VDRC #7723), *w;esg-GAL4,UAS-CFP, Su(H)-GFP;tubGal80*^*ts*^ *(esg*^*ts*^,*UAS-CFP,SuH-GFP)* (Lucy O’Brein), *UAS-relVP16* (Bloomington #36547), *UAS-RasN17* (Bloomington #4845), and *UAS-RasV12* (Bloomington #4847).

### Immunofluorescence

Intestines were dissected in PBS, fixed in 8% formaldehyde for 20 minutes, washed in PBS 0.2% Triton-X (PBST) then blocked in PBST with 3% BSA for 1hr at room temperature. Primary antibodies were incubated in PBST with BSA overnight at 4°C. The following day guts were washed in PBST then secondary antibody incubations were done in conjunction with DNA stain for 1 hour at room temperature in PBST with BSA, washed with PBST then again with PBS. Primary antibodies used: anti-prospero (1/100; Developmental Studies Hybridoma Bank (DSHB) MR1A), anti-armadillo (1/100;DSHB N2 7A1), chicken anti-GFP (1/2000; Invitrogen PA1-9533), anti-phospho-histone3 (1/1000; Millipore 06-570), anti-Delta (1/100; DSHB C594.9B), anti-pERK (1/1000; Millipore 05-797R). Secondary antibodies used: goat anti-chicken 488 (1/1000; Invitrogen A11039), goat anti-mouse 568 (1/1000; Invitrogen A11004), goat anti-rabbit 568 (1/1000; Invitrogen A11011), goat anti-mouse 647 (1/1000; Invitrogen A21235), and goat anti-rabbit 647 (1/1000; Invitrogen A21244). DNA stains used: Hoechst (1/1000; Molecular Probes H-3569). Apoptotic cells were detected in dissected guts using the TMR red In Situ Cell Death Detection Kit (Roche; 12156792910) following standard kit staining protocol. Briefly, guts were washed in PBS following secondary antibody then stained with 100μL of TUNEL solution for 1hr at 37°C then washed twice with PBS. Intestines were mounted on slides using Fluoromount (Sigma; F4680). For every experiment, images were obtained of the posterior midgut region (R4/5) of the intestine with a spinning disk confocal microscope (Quorum WaveFX). PH3+ cells were counted through the entire midgut.

### DSS and *Ecc15* treatments

DSS was prepared by dissolving DSS (Sigma 42867) in a PBS 5% sucrose solution, filter sterilized and kept in the freezer for up to two weeks. A 3% DSS solution was used for Figures 1-4 and a 5% DSS solution was used for Figure 5 and Figure 3E-G for a more robust proliferative response. *Ecc15* was prepped by streaking then incubating LB plates at 29°C overnight, then inoculating LB broth (Difco Luria Broth Base, Miller, 241420 supplemented with 4.75g NaCl per 500mL broth) with single colonies and grown with shaking at 29°C for ~20-24hr. The liquid culture was spun down at 1250g for 10min and the bacterial pellet was resuspended in residual LB and diluted 1:1 in PBS 5% sucrose. DSS or *Ecc15* vials were prepped by covering normal fly food with circular filter paper (Whatman, Grade 3, 23mm, 1003-323) and adding 150μL of the DSS, *Ecc15* or unchallenged (PBS 5% sucrose) solution. Flies were flipped daily onto fresh DSS, *Ecc15* or control solution for 48hr for all experiments except for DSS survival which was over the course of 10 days.

### Lifespan

For longevity, 30 virgin females per vial were raised at 29°C and dead flies were counted every 1-3 days and vials were flipped 3 times per week to fresh standard food. For DSS survival experiments flies were placed on fresh 10% DSS daily for the course of the survival experiment and deaths were counted daily.

### Data visualization and Statistical analysis

Figures were constructed using R (version 4.1.2) via R studio with easyggplot2 (version 1.0.0.9000) or ggplot2 (version 3.3.5). Statistical analysis was performed in R. Figures were assembled in Adobe Illustrator.

### Sample prep for single cell RNA sequencing

Preparation of single-cell intestinal suspension was made following previous methods (31, 44). Flies were raised for 10 days at 29°C then treated with 3% DSS or unchallenged solution for 48hrs. Batches of five *Drosophila* midguts were dissected at once then transferred to 1% BSA in DEPC treated PBS. Once 30 midguts were obtained for each condition they were transferred to a 1.5mL tube with 200μL of DEPC/PBS with 1mg/mL Elastase (Sigma, E0258) and chopped into pieces with small dissecting scissors. After mechanical disruption, tubes were incubated at 27°C for 40min with gentle pipetting every 10min. 22μL of 10%BSA in DEPC/PBS solution was added to stop the enzymatic disruption then cells were pelleted by spinning at 300g for 15min at 4°C. Cell pellet was resuspended in 200μL of 0.04% BSA in DEPC/PBS then filtered through a 70μM filter. Live cells were enriched using OptiPrep Density Gradient Medium (Sigma, D1556). Filtered cells were mixed with 444μL of 40% iodixanol (2:1 OptiPrep:0.04% BSA DEPC/PBS) then transferred to a 15mL tube. Another 5.36mL of 40% iodixanol was added and mixed. A 3mL layer of 22% iodixanol was added on top then an additional 0.5mL layer of 0.04% BAS in PBS/DEPC was added. Tubes were spun at 800g for 20min at 20°C then the top interface containing live cells (~500μL) was collected. Live cells were diluted with 1mL of 0.04% BSA in DEPC/PBS. Remaining iodixanol was removed by pelleting cells at 300g for 10min at 4°C and removing supernatant. Cell pellet was resuspended in remaining 0.04% BSA DEPC/PBS solution (~40μL) and cell counts and viability was determined using a hemocytometer. Cell viability was as follows: *ISC*^*ts*^*/+* unchallenged = 96%, *ISC*^*ts*^*/+* DSS = 95%, *ISC*^*ts*^/*relRNAi* unchallenged = 97%, *ISC*^*ts*^/*relRNAi* DSS = 86%. Libraries were generated using 10X Genomics Single-cell Transcriptome Library kit and sent to Novogene for sequencing.

### Bioinformatics

Raw sequencing data from Novogene was aligned to the *Drosophila* reference transcriptome (FlyBase, r6.30) using Cell Ranger v3.0 with the EYFP sequence appended to generate feature-barcode matrices. The resulting matrices were analyzed using Seurat (v4.1.0)(45, 46) in R. Cells with <200 or >3500 features and cells with >20% mitochondrial reads were removed to reduce number of low quality cells or doublets. Expression values were normalized and data clustering was performed at a resolution of 0.4 with 30 principal components. Clusters were identified using established markers and previous *Drosophila* intestine single-cell analysis (www.flyrnai.org/scRNA)(31, 44). GO term analysis was performed using Gorilla using unranked two list.

### Data Availability

Gene expression data is deposited on NCBI under the accession number PRJNA873108.

## SUPPLEMENTARY FIGURES

**Supplementary Figure 1.**
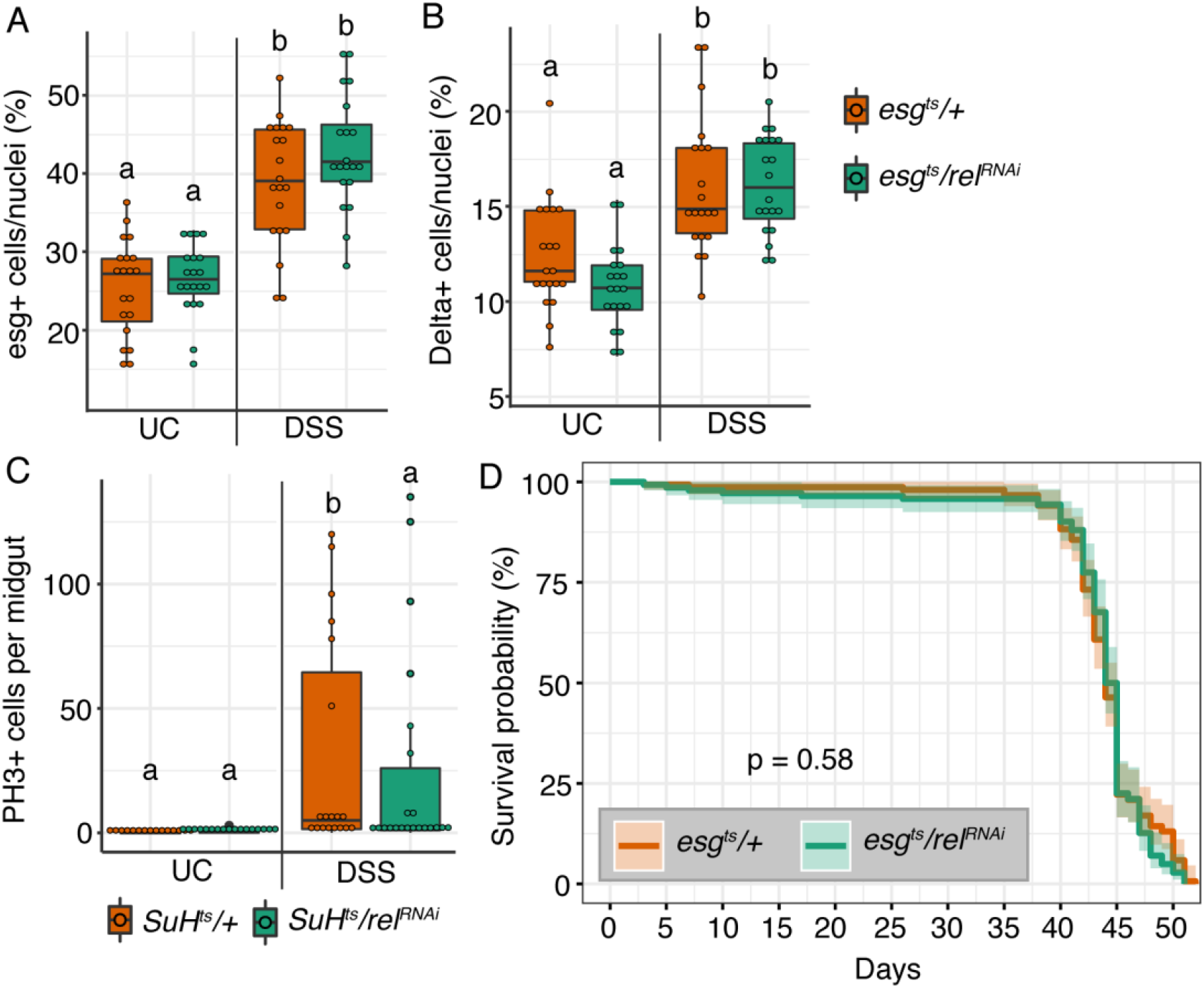
Progenitor-specific NF-κB does not alter progenitor numbers or fly survival upon damage. **(A)** Proportion of nuclei that are esg+ progenitors upon progenitor specific *rel* knockdown and DSS exposure. **(B)** Proportion of nuclei that are Delta+ ISCs upon progenitor specific *rel* knockdown and DSS exposure. **(C)** PH3+ cells per midgut upon *rel* knockdown in enteroblasts (*SuH*^*ts*^*/rel*^*RNAi*^) and DSS exposure. For A-C significance found using ANOVA followed by pairwise Tukey tests. Different letters denote significance at p < 0.05. **(D)** Lifespan analysis of *esg*^*ts*^*/+* and *esg*^*ts*^*/rel*^*RNAi*^ raised under normal conditions. Significance found using log rank test.

**Supplementary Figure 2.**
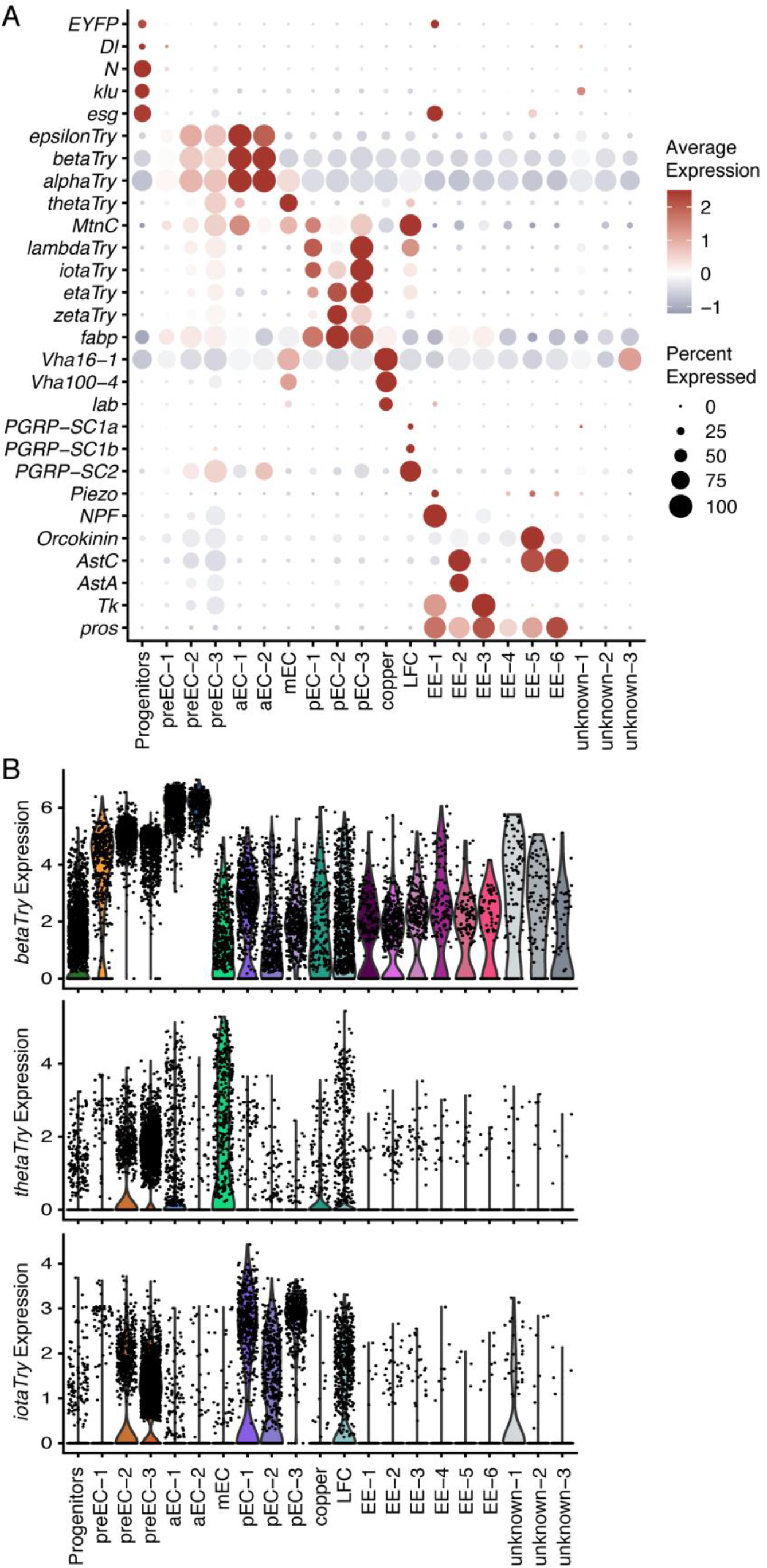
Premature enterocytes express anterior, middle and posterior enterocyte markers. **(A)** Dotplot from 4-way integrated single-cell sequencing data showing marker genes enriched in the identified cell clusters. Size of the dot represents the proportion of cells within the cluster that express that gene. Dot color represents the average expression level for that gene within the specified cluster. **(B)** Violin plots showing the expression (*thetaTry*), enterocyte of anterior (*betaTry*), middle and posterior (*iotaTry*) markers genes enriched in preECs. Each dot represents an individual cell.

**Supplementary Figure 3.**
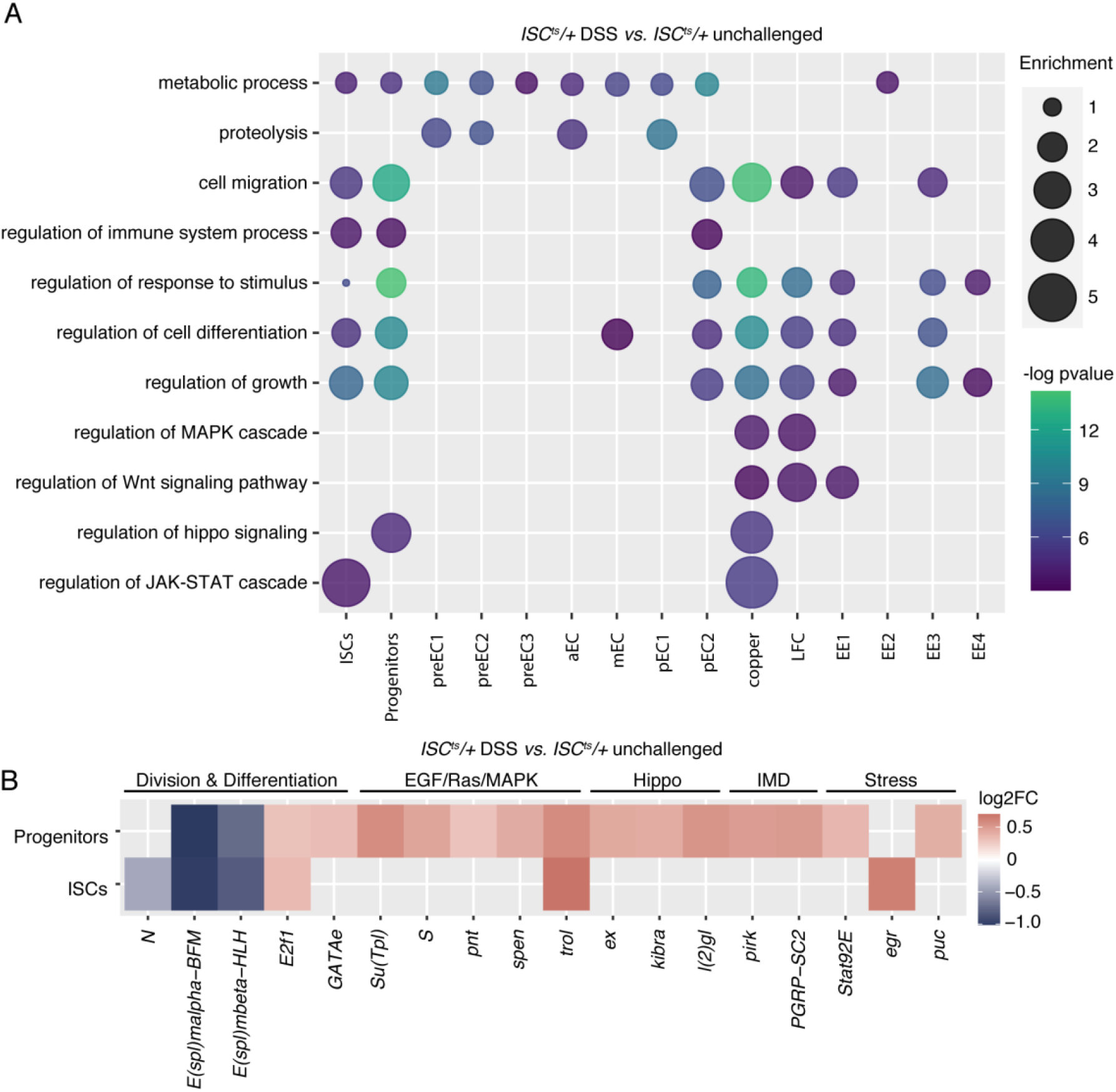
DSS exposure activates growth and stress pathways throughout the intestine. **(A)** GO term plot showing biological processes significantly altered in wild-type intestines upon exposure to DSS. Size of the bubble shows GO term enrichment score and color shows −log pvalue. **(B)** Genes differentially expressed in Progenitors and EYFP+ Progenitors (ISCs) in DSS treated *ISC*^*ts*^*/+* intestines compared to unchallenged *ISC*^*ts*^*/+*. All genes shown are p < 0.05.

**Supplementary Figure 4.**
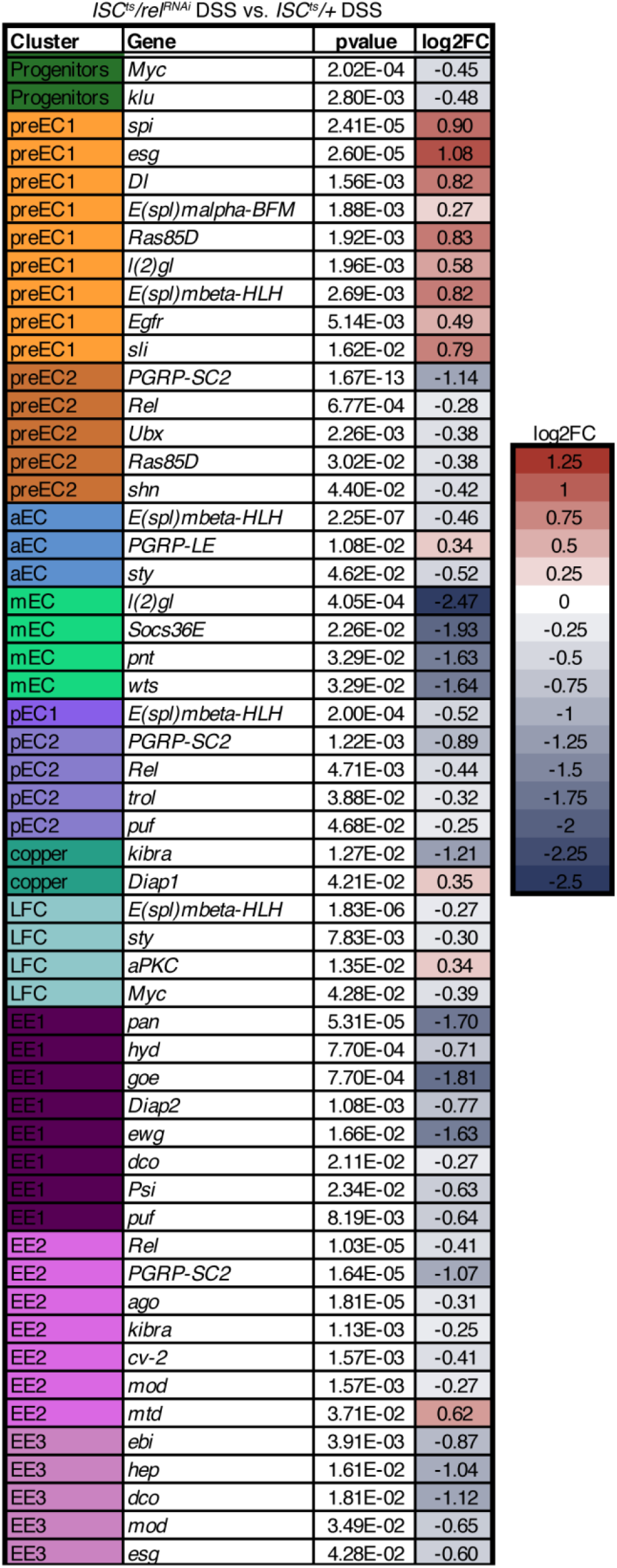
ISC-specific Relish alters immune, growth and stress genes throughout the epithelium in response to DSS. Comparison of DSS treated *ISC*^*ts*^*/rel*^*RNAi*^ intestines to DSS treated *ISC*^*ts*^*/+*. Differentially expressed genes across cell clusters involved in IMD pathway, ISC division, differentiation, stress and homeostasis. All genes shown are p < 0.05.

**Supplementary Figure 5.**
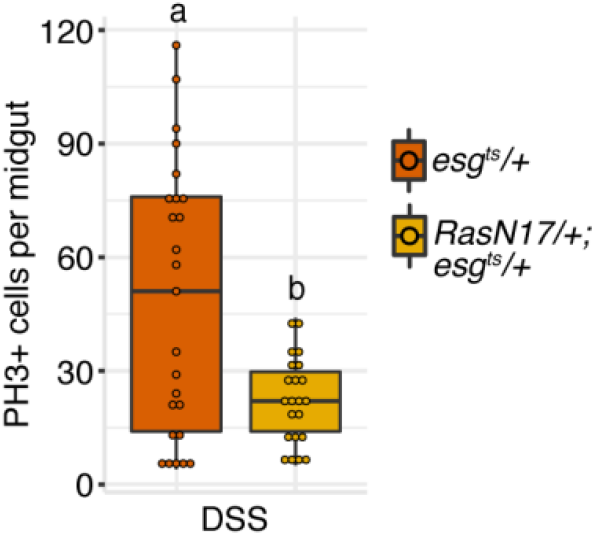
Progenitor-specific Ras promotes ISC divisions in response to DSS. PH3+ cells per midgut after progenitor-specific Ras inactivation (*RasN17/+;esg*^*ts*^*/+)*. Significance found using Students t test. Different letters denote significance at p <0.05.

## Notes

### Competing Interest Statement

The authors have declared no competing interest.

